# Integrated Hfq-interacting RNAome and transcriptomic analysis reveals complex regulatory networks of nitrogen fixation in root-associated *Pseudomonas stutzeri* A1501

**DOI:** 10.1101/2023.12.18.572192

**Authors:** Fanyang Lv, Yuhua Zhan, Haicao Feng, Wenyue Sun, Changyan Yin, Yueyue Han, Yahui Shao, Wei Xue, Shanshan Jiang, Yiyuan Ma, Haonan Hu, Wei Jinfeng, Yongliang Yan, Min Lin

**Author notes:** Correspondence (Y.Y.) and (M.L.).

## Abstract

The RNA chaperone Hfq acts as a global regulator of numerous biological processes, such as carbon/nitrogen metabolism and environmental adaptation in plant-associated diazotrophs; however, its target RNAs and the mechanisms underlying nitrogen fixation remain largely unknown. Here, we used enhanced UV cross-linking immunoprecipitation coupled with high-throughput sequencing (eCLIP-seq) to identify hundreds of Hfq-binding RNAs probably involved in nitrogen fixation, carbon substrate utilization, biofilm formation, and other functions. Collectively, these processes endow strain A1501 with the requisite capabilities to thrive in the highly competitive rhizosphere. Our findings revealed a previously uncharted landscape of Hfq target genes. Notable among these is *nifM*, encoding an isomerase necessary for nitrogenase reductase solubility; *amtB,* encoding an ammonium transporter; *oprB,* encoding a carbohydrate porin; and *cheZ,* encoding a chemotaxis protein. Furthermore, we identified more than one hundred genes of unknown function, which expands the potential direct regulatory targets of Hfq in diazotrophs. Our data showed that Hfq directly interacts with regulatory proteins (RsmA, AlgU, NifA), regulatory ncRNA RsmY, and other potential targets, thus revealing the mechanistic links in nitrogen fixation and other metabolic pathways.

**IMPORTANCE:** Numerous experimental approaches often face challenges in distinguishing between direct and indirect effects of Hfq-mediated regulation. New technologies based on high-throughput sequencing are increasingly providing insight into the global regulation of Hfq in gene expression. Here, enhanced UV cross-linking immunoprecipitation coupled with high-throughput sequencing (eCLIP-seq) was employed to identify the Hfq-binding sites and potential targets in the root-associated *P. stutzeri* A1501, and identify hundreds of novel Hfq-binding RNAs that are predicted to be involved in metabolism, environmental adaptation, and nitrogen fixation. In particular, we have shown that Hfq interactions with various regulatory proteins and their potential targets at both the protein and RNA levels. This study not only enhances our understanding of Hfq regulation but, importantly, also provides a framework for addressing integrated regulatory network underlying root-associated nitrogen fixation.

## INTRODUCTION

Hfq is a ubiquitous RNA-binding protein (RBP), originally identified in *Escherichia coli* as a host factor essential for the in vitro synthesis of RNA bacteriophage Qβ-RNA(1), and has now been widely recognized as an RNA matchmaker (2). Due to its ability to bind a diverse array of RNAs, Hfq has been implicated in a multitude of complex phenotypes, including motility, carbon metabolism, host colonization, nitrogen fixation, fitness, and virulence of bacteria (5, 6, 7, 8, 9). The primary mode of action of Hfq is through acceleration of sRNA-mRNA annealing and subsequent RNA stabilization or degradation though alternative regulatory mechanisms(2, 3, 4). Furthermore, Hfq has been shown to independently interact with either ncRNAs or mRNAs, although it may not necessarily promote interaction between these molecules; for example, Hfq directly binds to sites containing the (AAN)_n_ motif in the 5’UTR of mRNAs, affecting gene translation (10, 11). Moreover, Hfq-dependent ncRNAs undergo rapid degradation when they are not bound to Hfq, involving a process wherein RNase E and PNPase function as active degradative enzymes (12, 13). Additionally, Hfq can modulate mRNA stability, including that of its own transcript, by directly binding and remodeling the mRNA molecule independently of ncRNA(10, 14, 15). Despite the considerable interest in establishing a universal role for Hfq, the in vivo global binding preferences of this protein remain unknown.

Numerous experimental approaches have also been developed to facilitate the identification of Hfq targets in bacteria (17, 18). However, these methods often face challenges in distinguishing between direct and indirect effects of Hfq-mediated regulation. New technologies based on high-throughput sequencing are increasingly providing insight into the global regulation of Hfq in gene expression(4). In particular, enhanced UV cross-linking immunoprecipitation coupled with high-throughput sequencing (eCLIP-seq) is a variant of CLIP-seq with improved sensitivity and specificity and has been widely used to unravel the RBP interactome (19,20,21). Several coimmunoprecipitation studies have revealed that Hfq binds hundreds of mRNAs and ncRNAs in the model pathogen *Salmonella typhimurium* (22), the most frequent opportunistic biofilm-forming bacterium *Pseudomonas aeruginosa* (23), and the nitrogen-fixing legume symbiotic species *Sinorhizobium meliloti* (24). These findings expand our understanding of the potential direct regulatory targets of Hfq in bacteria.

*Pseudomonas* are ubiquitous bacteria that can live in a wide range of environments, such as soil ecosystems and associations with plants and animals (25). *P. stutzeri* A1501 stands out as a nitrogen-fixing strain within the Pseudomonas genus (26). The A1501 strain harbors the *glnB/glnK/amtB/ntrBC/RpoN* genes, constituting a fundamental nitrogen regulatory system that governs nitrogen and carbon mechanisms and is embedded in the core genome. Evolutionary processes have endowed A1501 with a *nif*-specific regulatory system (*nifLA*) through horizontal gene transfer from a diazotrophic ancestor (26). Furthermore, a multifunctional regulatory RNA, NfiS, has been recruited by *nifK* mRNA, serving as a novel activator to optimize the nitrogen fixation process in response to specific environmental cues (27). Thus, the regulatory network controlling nitrogen fixation in A1501 likely arises from regulatory systems of different evolutionary origins. In addition, A1501 is isolated from the rice rhizosphere and exists either in a free-living lifestyle in the soil or in root association with host plants. The successful transition from the free-living state to rhizosphere colonization involves massive reprogramming of gene expression by both regulatory proteins and ncRNAs, such as the regulatory network involving RpoS/RsmA/RsmZY for nitrogen-fixing biofilm formation and the Hfq/Crc/CrcZY regulatory system underlying hierarchical carbon substrate utilization (9, 28). Random chemical mutagenesis screenings in *Azorhizobium caulinodans* identified Hfq as an activator for translation of NifA, one of the master regulators of nitrogen fixation(29). Additionally, Hfq was also found to influence an array of *S. meliloti* symbiotic traits, including competitiveness for infection, nodule development, intracellular survival of bacteroids, and efficiency of the nitrogen fixation process(24, 30). Likewise, Hfq has been shown to exert a global effect on A1501. The *hfq* mutant of A1501 has also been shown to be severely impaired in nitrogen fixation, carbon substrate utilization, exopolysaccharide biosynthesis, and root colonization(9, 27). Consistent with observations in other bacteria, Hfq in A1501 is involved in a broader regulatory spectrum, affecting diverse mechanisms of action, including repressing the translation of substrate-specific catabolic genes, activating both the nitrogenase gene *nifH* at the posttranscriptional level and an exopolysaccharide gene cluster at the transcriptional level, particularly affecting the stability of regulatory ncRNAs associated with environmental stresses or induced under nitrogen fixation conditions(9). Obviously, Hfq acts as a global regulator of numerous biological processes in A1501; however, detailed information pertaining to its regulatory targets was available for only a few instances. New approaches are needed to fully elucidate Hfq-mediated mechanisms underlying nitrogen fixation and plant–microbe interactions.

We previously reported a global transcriptional profiling analysis of nitrogen fixation and ammonium repression in A1501 and identified a total of 95 genes as part of the core subset of the regulon induced specifically under nitrogen fixation conditions (31). This subset includes not only the *nif* genes (20%), which are directly involved in the synthesis, maturation, and function of nitrogenase but also other core genes encoding global regulators, transport proteins and metabolic enzymes (39%), as well as proteins involved in energy production and conversion (16%). More recently, a total of 53 ncRNAs were detected under nitrogen fixation conditions, 17 of which are upregulated under nitrogen fixation conditions but were rapidly downregulated after 10 min of ammonium shock (32). Notably, a substantial proportion of genes encoding both proteins and ncRNAs involved in gene expression and energy metabolism are induced under nitrogen fixation conditions, aligning with the well-known fact that biological nitrogen fixation is a highly regulated and energy-dependent process. To ascertain whether the expression of nitrogen fixation-inducible genes is directly regulated by Hfq, we investigated the Hfq regulon in *P. stutzeri* A1501 by using genome-wide RNA-seq analyses of *hfq* mutants and coimmunoprecipitation with tagged Hfq. RNA sequencing (RNA-seq) was employed to identify the impact of Hfq on RNA abundance and expression, while qRT‒PCR was utilized to measure the half-life of ncRNAs/mRNAs in the wild-type (WT) and Δ*hfq* strains. This multiple omics approach not only expands the potential direct regulatory targets for Hfq but also promises to yield novel insights into the global regulatory mechanisms orchestrated by Hfq in the context of nitrogen fixation.

## RESULTS

### Genome-wide profiling of Hfq-binding RNAs and their expression under nitrogen fixation conditions

To investigate the interactions of Hfq with target RNAs under nitrogen fixation conditions, we used cells of *P. stutzeri* A1501 chromosomally encoded FLAG epitope-tagged Hfq variant (Hfq^FLAG^) to comprehensively identify Hfq-bound transcripts. Wild-type cells that synthesize untagged Hfq served as a control to assess unspecific RNA binding to either the magnetic beads used for CoIP or the FLAG tag itself. As shown in Fig. S1, cells were grown under nitrogen fixation conditions and exposed to UV to allow cross-linking of RNAs to Hfq. Following cell lysis, Hfq–RNA complexes from crosslinked and control samples were coimmunoprecipitated using anti-FLAG antibody. Exposed regions of the RNAs were trimmed by RNases, the Hfq protein was digested, neighboring RNAs were ligated, and the resulting RNA was subjected to high-throughput paired-end sequencing. Upon normalization, the distribution of the sequencing reads from Hfq CoIP and control samples was visualized using the integrated genome browser (Fig. 1A). Mapping of the sequenced reads was performed on the *P. stutzeri* A150 genome, and hundreds of Hfq-binding reads were identified as peaks using the tool Clipper (33) (Table S1).

**Fig. 1:**
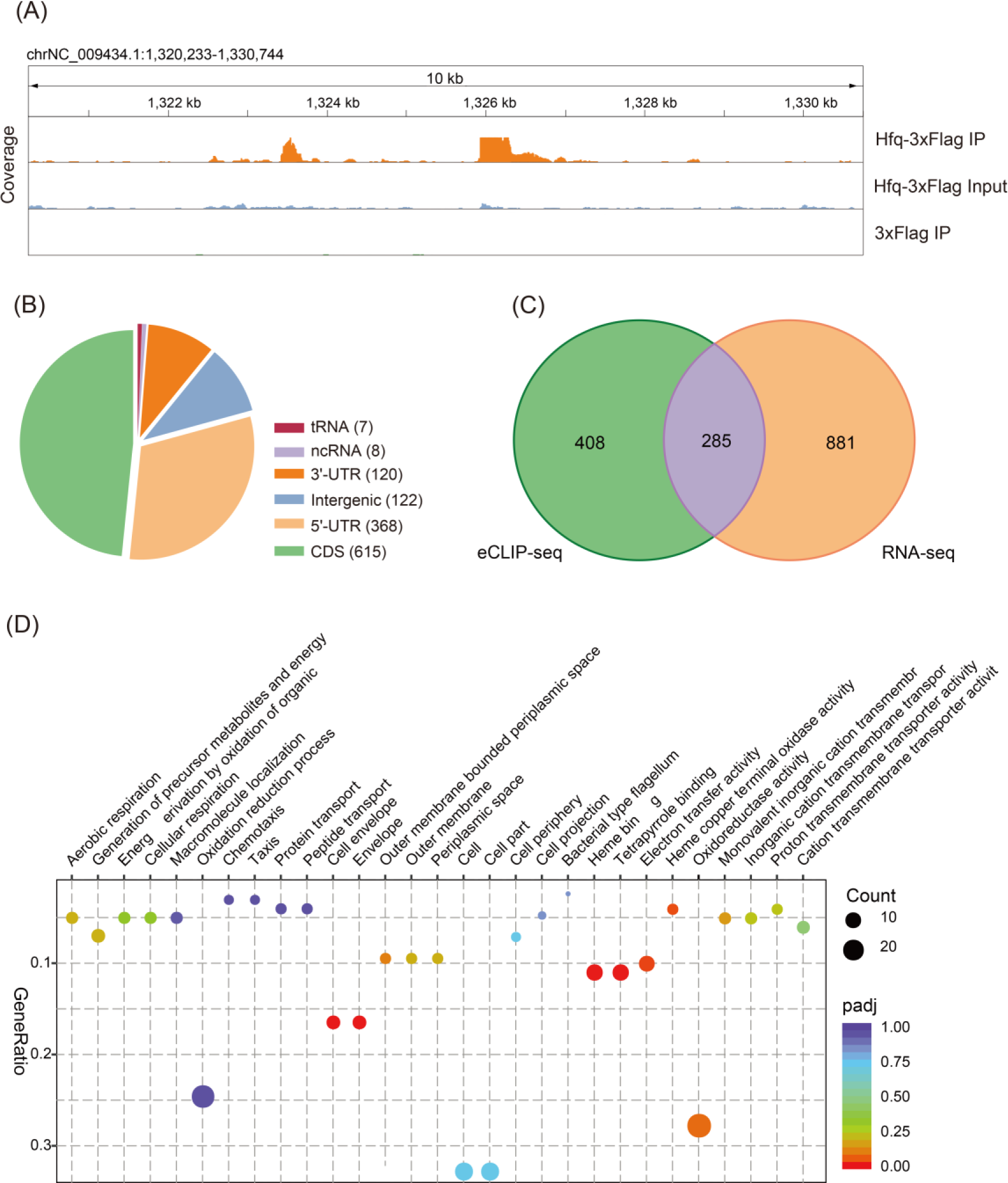
Global patterns of Hfq-binding targets. (A). IGV (Integrative Genomic Viewer) snapshot of genomic regions with eCLIP-seq data. Reads from eCLIP-seq with A1501 Hfq-3xFlag IP cells compared to eCLIP-seq Input cells and 3xFlag cells. (B). Localization of the Hfq eCLIP-seq peaks relative to the predicted CDS, ncRNA, 5’UTR, and 3’UTR. (C). Venn diagram depicting genes with significant expression changes according to the different transcriptomic approaches. eCLIP-seq peaks without associated annotations are not shown. (D). GO enrichment analysis of differentially expressed genes between the WT and Δ*hfq* strains according to both RNA-seq and Hfq eCLIP-seq peak data. ‘Gene ratio’ shows the ratio of genes related to the GO term to the total number of differentially expressed genes annotated with the given GO term of biological processes and molecular functions identified using DAVID to be enriched.

Based on their genomic context with respect to a minimal transcription unit model, 987 peaks in the experimental library with respect to the control were regarded as Hfq-bound and were further cataloged as shown in Fig. 1B and Table S1: (1) 615 mRNAs (62% of the peaks), if matched the sense strand of protein-coding regions, including their experimentally determined or virtual 5’/3’untranslated regions; (2) eight previously recognized ncRNAs, those corresponding to several peaks for remaining unannotated intergenic regions, and seven tRNA coding genes; and (3) 122 Hfq-binding RNAs (12%), whose function are unknown.

We further used RNA-seq to analyze differences in RNA abundance and expression between the WT and Δ*hfq* strains (Fig. S1). To globally assess the Hfq regulon, we compared the transcriptome of wild-type cells with that of the *hfq* mutant strain, both grown under nitrogen fixation conditions. The differentially accumulated transcripts were identified by setting a *p* value of 0.05 and a log^(FoldChange [FC])^ threshold ≥1.0 or ≤ −1.0. A total of 1,093 genes whose RNA abundance was significantly altered (Table S2) accounted for approximately 26% of *P. stutzeri* A1501 annotated open reading frames compared with those of the WT. The abundance of approximately 53% of the differentially accumulated transcripts was decreased in Δ*hfq,* including *pslA* and *nifH*, which had been reported previously (9), whereas the abundance of 47% of these transcripts was increased. Among the upregulated genes, translation initiation of *benR*, *estA*, and *mupP* was previously reported to be directly controlled by Hfq (34). We also analyzed the effects of Hfq on RNA abundance for nonprotein-coding genes, which detected 110 ncRNAs showing differential expression (Table S3); 103 out of 110 were downregulated in Δ*hfq* compared with the WT, suggesting a key role of Hfq in the positive regulation of gene expression.

Gene Ontology (GO) enrichment analysis indicated that the genes associated with the eCLIP-seq peaks were enriched in a large array of biological processes (51%), molecular functions (42%), and cellular components (7%), whose expression was significantly affected according to RNA-seq (Fig. S3A). Transcriptome sequencing data from the *P. stutzeri* A1501 strains cultured under the same conditions as another control dataset (Table S2) showed that non-Hfq-binding genes were enriched in flagellar assembly, bacterial secretion system, and chemotaxis (Table S2, Fig. S3B). The overlap of genes identified using the two transcriptomic approaches is depicted in Fig. 1C and 1D, indicating that Hfq influences most genes critical for the growth of *P. stutzeri* A1501, which supports the notion of its global and extensive regulatory role. Overall, our CLIP-seq and transcriptome approach will provide new insights into Hfq-mediated global regulation under nitrogen fixation conditions.

### Enrichment analysis of Hfq-dependent RNAs involved in nitrogen fixation and other associated metabolic pathways

Biological nitrogen fixation is tightly regulated at both the transcriptional and posttranslational levels in response to the availability of fixed nitrogen. It is also a highly energy-dependent process requiring large amounts of carbon and energy sources (35). We previously identified a total of 95 protein-coding genes and 17 ncRNA-coding genes induced specifically under nitrogen fixation conditions(31). In this study, the enrichment analysis revealed that Hfq directly regulates genes involved in ammonium uptake and metabolism and various functions related to nitrogen fixation (Table S1, Fig. 2A). We identified an eCLIP-seq peak corresponding to the CDS of *glnA*, encoding a glutamine synthetase, which lies at the heart of the nitrogen assimilation network (36) and was decreased in Δ*hfq* nearly 2-fold in Δ*hfq* compared with the WT (Table S2). We further confirmed that Hfq repressed *glnA* expression by using qRT‒PCR and *glnA::lacZ* translation fusions (Fig. 2B, 2C). Thus, Hfq binding appears to activate glutamine synthetase expression. Additionally, we found an eCLIP-seq peak corresponding to the 5’UTR of *amtB* (Table S1), encoding the high-affinity ammonium transporter, in an operon together with a second gene (*glnK*) encoding a small signal transduction protein, GlnK, which acts as a sensor of the cellular nitrogen status in prokaryotes (37). Both *amtB* and *glnK* were upregulated approximately 2-fold in Δ*hfq* (Tables S1 and S2).

**Fig. 2:**
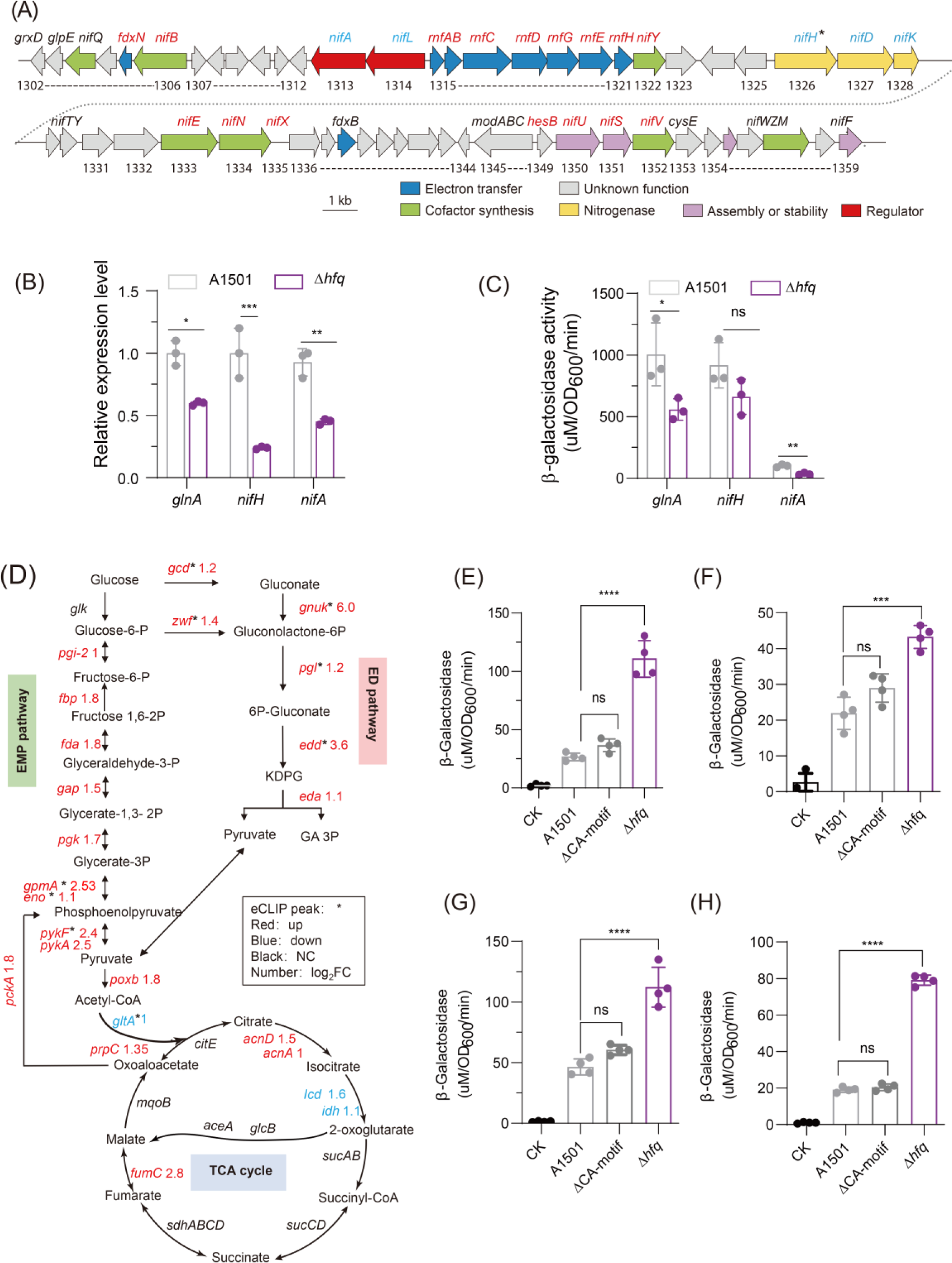
Effects of Hfq on the expression of genes involved in nitrogen fixation and central carbon metabolism. (A). Effect of Hfq on nitrogen fixation island genes. (B, C) Relative expression levels of *glnA*, *nifH* and *nifA* (B) and β-galactosidase activity (C) in the WT versus the *hfq* deletion mutant. (D). Effects of Hfq on the expression of genes involved in central carbon metabolism in *P. stutzeri* A1501. (E, F, G, H). The bars depict the β-galactosidase levels conferred by the chromosomally integrated translational *zwf*::*lacZ* (E) *gcd*::*lacZ* (F), *oprB*::*lacZ* (G), and *gtsA*::*lacZ* (H) fusion constructs expressed in WT and Δ*hfq*, respectively. The CA motif indicates the sequence containing the AANAANAA motif in the 5’UTR of the above genes. The asterisks indicate that a Hfq eCLIP-seq peak is associated with the gene. Red indicates repression by Hfq, blue indicates activation, and black indicates no change (NC) in gene expression. The number indicates the fold change (log_2_FC). Asterisks indicate statistical significance determined by one-way ANOVA with Tukey’s post hoc test: ****: *p* ≤ 0.0001; ***: *p* ≤ 0.001; ** *p* ≤ 0.01, * *p* ≤ 0.05; ns: not significant.

Many nitrogen fixation-related genes that are regulated by Hfq are controlled via the master regulator of nitrogen fixation (*nif*) gene NifA (a member of the bacterial enhance-binding family), the activity of which is controlled by a partner protein, NifL(29). Hfq mutation led to a significant decrease in the expression of both *nifA* and *nifL* under nitrogen fixation conditions (Fig. 2A, 2B, 2C). qRT‒PCR and β-galactosidase activity further confirmed that the absence of Hfq resulted in a significant decrease in the expression of *nifA* (Fig. 2B, 2C). The CDS of *nifH* (encoding the iron-containing electron transfer protein), which is regulated by NifA, exhibited an Hfq eCLIP-seq peak (Fig. 2A) and was downregulated approximately 4-fold, as revealed by qRT‒PCR and *nifH::lacZ* translational infusion in the absence of Hfq (Fig. 2 B, C), suggesting a positive effect of Hfq on *nifH* expression

We also identified eCLIP-seq peaks corresponding to regions in the 5’UTR of the mRNAs that were repressed in the presence of Hfq, such as *zwf* (encoding a glucose-6-phosphate-1-dehydrogenase), *gcd* (encoding glucose dehydrogenase), and *edd* (encoding 6-phosphogluconated dehydratase), which were upregulated in Δ*hfq* compared with the WT strain (Fig. 2D; Table S2). The β-galactosidase activity was further conferred by the translated *zwf::lacZ* and *gcd::lacZ* fusion constructs in WT and Δ*hfq* strains grown in medium K containing glucose. Unlike in the WT, *zwf::lacZ* and *gcd::lacZ* translation was significantly increased in the Δ*hfq* strain (Fig. 2E, 2F), whereas mutation of the presumptive binding site (AANAANAAN (CA-motif) to TCAGTAGC in the 5’ untranslated region of the target genes) led to a significant decrease in β-galactosidase activity (Fig. 2E, 2F). Thus, Hfq appears to repress the EMP and ED pathways by binding to *zwf, gcd* and other mRNA 5’UTRs (Fig. 2D), reducing the expression of the genes. Additional central carbon metabolism-related genes exhibited Hfq eCLIP-seq peaks and may be directly regulated by Hfq (Fig. 2D).

Genes involved in the transport of carbon sources were enriched in our eCLIP-seq data (Table S1) and Hfq-mediated transcriptome profile (Table S2), which included many gene clusters encoding components of the ABC transport system. For example, Hfq represses genes involved in the transport of glucose (*oprB*, *gtsABC*), benzoate (*benFK*), mannitol (*mltKEG*), and cystine (*tcyJ/L*) and activates genes encoding proteins with branched-chain amino acids (*braF*, *brnQ*) and choline (*PST4102*) (Table S2). We further confirmed that Hfq repressed *oprB* and *gtsA* expression by using chromosomal *oprB::lacZ* and *gtsA::lacZ* translation fusions in K media containing glucose (Fig. 2G, 2H). These carbon sources may be important for the bacterial growth of rice roots in certain ecological niches. For instance, glucose, some branched-chain amino acids, formate, and betaine support the growth of *Pseudomonas* in the rice root rhizosphere or improve the resistance of rice (38). Overall, Hfq directly regulates the expression of (> 200) genes that encode transporters of carbon sources and other nutrients and whose expression was significantly altered. These data further support that Hfq functions as a key player involved in the regulation of nitrogen fixation and other associated metabolic pathways.

### Identification of Hfq-dependent RNAs involved in chemotaxis and biofilm formation

Chemotaxis in bacteria is controlled by regulating the direction of flagellar rotation, which allows bacteria to move in or against chemical concentration gradients and facilitates the colonization of more favorable ecological niches (39). Hfq has been shown to be involved in motility and chemotaxis in many bacterial species (40). The deletion of *hfq* led to a significant impairment in motility compared with the WT, but this decrease was eliminated by genetic complementation with a plasmid carrying the WT *hfq* gene (Fig. 3A). Hfq affects the expression of numerous genes involved in flagellum-driven chemotaxis in the A1501 strain, which is reflected by the results from the enrichment analysis shown in Tables S1 and S2. Hfq activates genes including *cheB* (ending a receptor-specific methylesterase, which contains a response regulator module that is activated when it is phosphorylated by CheA-P), *cheR* (a methyltransferase that is also involved in the switching mechanism), and *cheZ* (a specific phosphatase to *cheY*). Many of the effects appear to represent indirect changes in expression and are not associated with the eCLIP-seq peak, except CheZ. qRT‒PCR showed that *cheZ* and *cheR* expression was decreased in Δ*hfq* by approximately 2-fold, and the β-galactosidase activity of *cheZ::lacZ* was significantly reduced, whereas *cheR::lacZ* translational fusion showed no significant change, which may be due to an indirect effect (Fig. 3B, 3C). In addition, *flic*, *flgE*, and *flgK* were downregulated in Δ*hfq.* Hfq also repressed the expression of the *fliG*, *fliM*, *motA*, and *motB* genes, which were all upregulated in Δ*hfq* compared with WT (Table S2, Fig. 3B, 3C). Overall, the absence of Hfq switches cells to a clockwise rotation, which results in cell tumbling and directional changes.

**Fig. 3:**
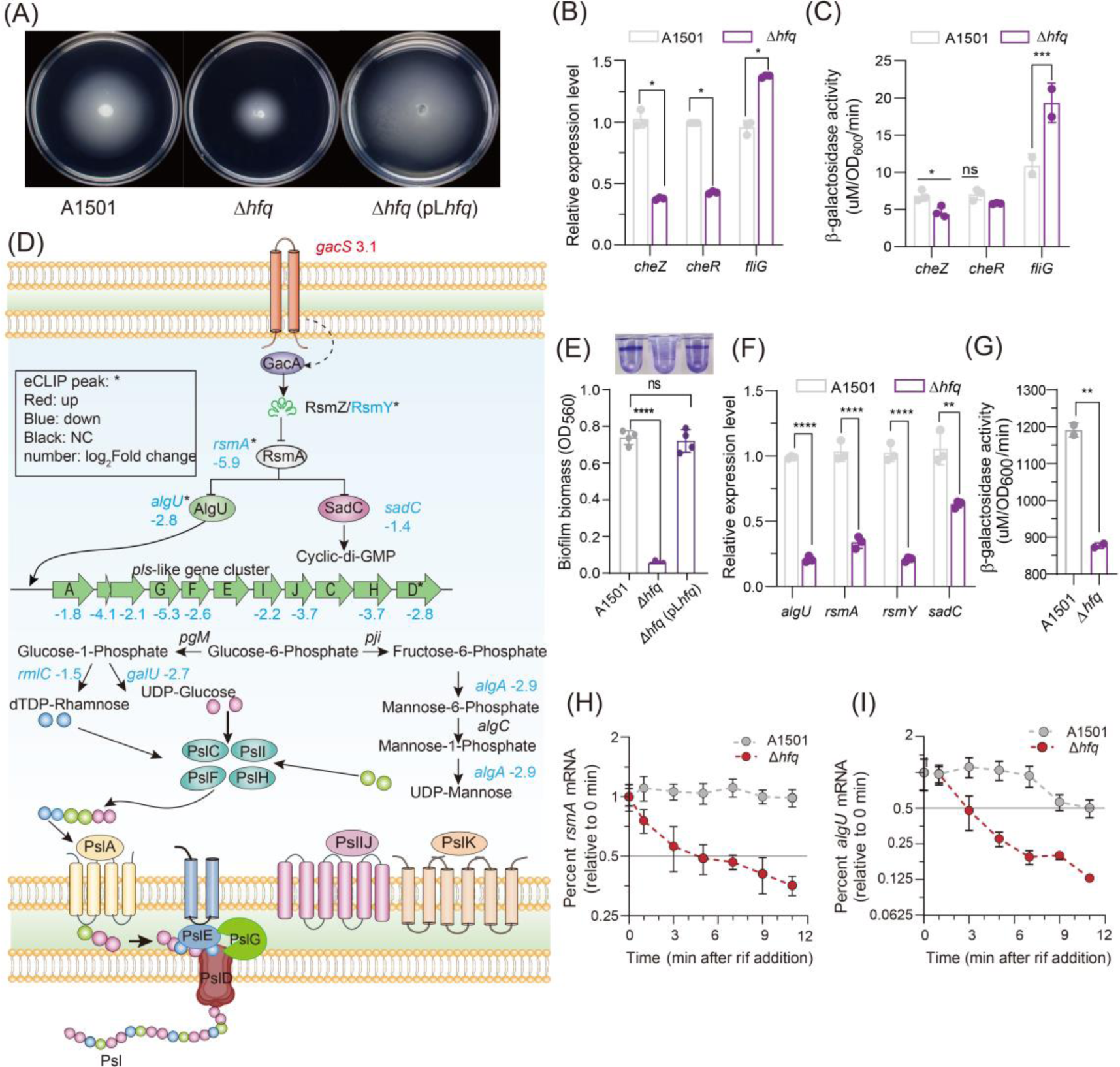
Effect of Hfq on the expression of genes involved in chemotaxis Mechanism and biofilm formation. (A). Phenotype of the chemotaxis ability of WT and Δ*hfq*. (B, C) Relative expression levels of *cheZ*, *cheR* and *fliG* (B) and β-galactosidase activity (C) in the WT and Δ*hfq*. (D). Summary of the effects of Hfq on biofilm formation and Psl production-related genes that were differentially expressed in the *hfq* mutant compared with the WT. (E). Effect of Hfq on biofilm formation after 48 h inoculation and comparison of the biofilm biomass obtained with the WT, *hfq* deletion mutant, and complemented strains. (F). Relative expression levels of *algU*, *rsmA*, *rsmY*, and *sadC* in the WT versus the *hfq* deletion mutant. (G). β-galactosidase activity of *rsm*A in the WT and Δ*hfq*. (H, I) Analysis of the *rsmA* (H) and *algU* (I) mRNA half-life in the WT and *hfq* mutant strains. Rifampicin (400 μg/mg) was added at time 0 min. The error bars show the standard deviations of the means of three independent experiments. Asterisks indicate that a Hfq eCLIP-seq peak is associated with the gene. Red indicates repression by Hfq, blue indicates activation, and black indicates no change (NC) in gene expression. The number indicates the fold change (log_2_FC). Asterisks indicate statistical significance determined by one-way (c) and two-way ANOVA with Tukey’s post hoc test: \*\*\*\**p* ≤ 0.0001, ***: *p* ≤ 0.001, \*\**p* ≤ 0.01, * *p*≤ 0.05, ns: not significant.

The deletion of *hfq* significantly decreased the biofilm formation ability compared with that of the WT (Fig. 3E). Hfq affects the expression of numerous genes involved in biofilm formation, as reflected by the results from the enrichment analysis shown Fig. 3D. Many related genes that are regulated by Hfq are controlled via RsmA, which binds to the 5’UTR of multiple mRNAs, enhancing bacterial motility while repressing the production of Psl (41). The 5’UTR of *rsmA* exhibited an Hfq eCLIP-seq peak in our study (Fig. 3D; Table S1). We further measured the half-life of the *rsmA* transcript, which was approximately 5 min in Δ*hfq*, whereas no changes were observed in the WT within 9 min (Fig. 3H). qRT‒PCR and β-galactosidase activity further confirmed that the absence of Hfq results in a significant decrease in the expression of *rsmA,* suggesting that Hfq directly binds *rsmA* mRNA and has a positive effect on its stability (Fig. 3F, 3G).

Following eCLIP-seq and RNA-seq transcriptomic analysis, qRT‒PCR was further used to measure the relative mRNA abundance of *algU*, RsmY, and *sadC*, which were significantly decreased in Δ*hfq* (Fig. 3F). AlgU (also known as AlgT and RpoE) functions as a sigma factor that regulates translation from a nonmucoid to mucoid state in *P. aeruginosa* and directly activates the transcription of *pslA* in *P. stutzeri* A1501 (42), which was downregulated nearly 2.8-fold (Fig. 3F). An eCLIP-seq peak associated with the 5’UTR *algU* mRNA was significantly enriched (Fig. 3D). We further measured the half-life of *algU* mRNA in the presence of rifampicin, which was 11 min in the WT strain but decreased to 3 min in the Δ*hfq* (Fig. 3I), indicating that Hfq exerted a positive effect on the stability of the *algU* mRNA. Hfq also activated genes involved in exopolysaccharide production, and deletion of *hfq* resulted in a significant decrease in the expression of all *psl*-like cluster genes (Fig. 3D), suggesting that indirect Hfq-mediated regulation occurred because the mRNAs of these genes did not exhibit an eCLIP-seq peak. Collectively, these genetic downregulation events explain the biofilm formation defect of the Δ*hfq* mutant.

### Hfq interactions with other global regulators and their integrated networks underlying nitrogen fixation

Consistent with known Hfq regulatory effects, numerous sequences associated with eCLIP-seq peaks overlapping 5’UTRs were associated with posttranscriptional regulation, primarily via negative effects on gene expression. In addition, sequences associated with eCLIP-seq peaks spanning the initiation sites of CDSs frequently exhibited a repressive profile.

Given that a substantial number of transcriptional changes manifest in *hfq*, independent of their association with eCLIP-seq peaks (Fig. 1C), it is reasonable to infer that these changes may be orchestrated by alternative regulatory elements, with a specific focus on sigma factors or transcriptional regulators. Indeed, the eCLIP-seq data revealed an enrichment of 30 transcriptional regulators and eight sigma factors (Table S1, Fig. S5A), and their Hfq-regulated status was further confirmed through qRT‒PCR and β-galactosidase activity assays in Δ*hfq* compared with WT (Fig. 4B, 4C, and Table S2). For instance, the mRNA *rpoS*, which exerts a negative regulatory effect on nitrogen-fixing biofilm formation via RsmZ (43), was found to be Hfq-bound and downregulated in Δ*hfq* compared with the WT (Fig. 4B-C), suggesting that Hfq has a positive effect on the stability and expression of *rpoS* mRNA. This possibility was checked by measuring the half-life of the *rpoS* transcript, which was approximately 11 min in the WT and reduced to 3 min in Δ*hfq* (Fig. 4E). Members of the ferric uptake regulator (Fur) protein family serve as bacterial transcriptional repressors for governing iron uptake and storage in response to iron availability, thereby playing a crucial role in maintaining iron homeostasis (44). Moreover, the ferric uptake regulator Fur mRNA, which is known to regulate the expression of the intracellular iron transport system in A1501(45), exhibited an Hfq eCLIP-seq peak, while its expression and half-life were decreased in Δ*hfq* (Fig. 4B-D). We also measured its half-life, which was 7 min in the WT but decreased to 5 min in Δ*hfq*, indicating that Hfq exerted a positive effect on the stability of *fur* mRNA (Fig. 4D).

**Fig. 4:**
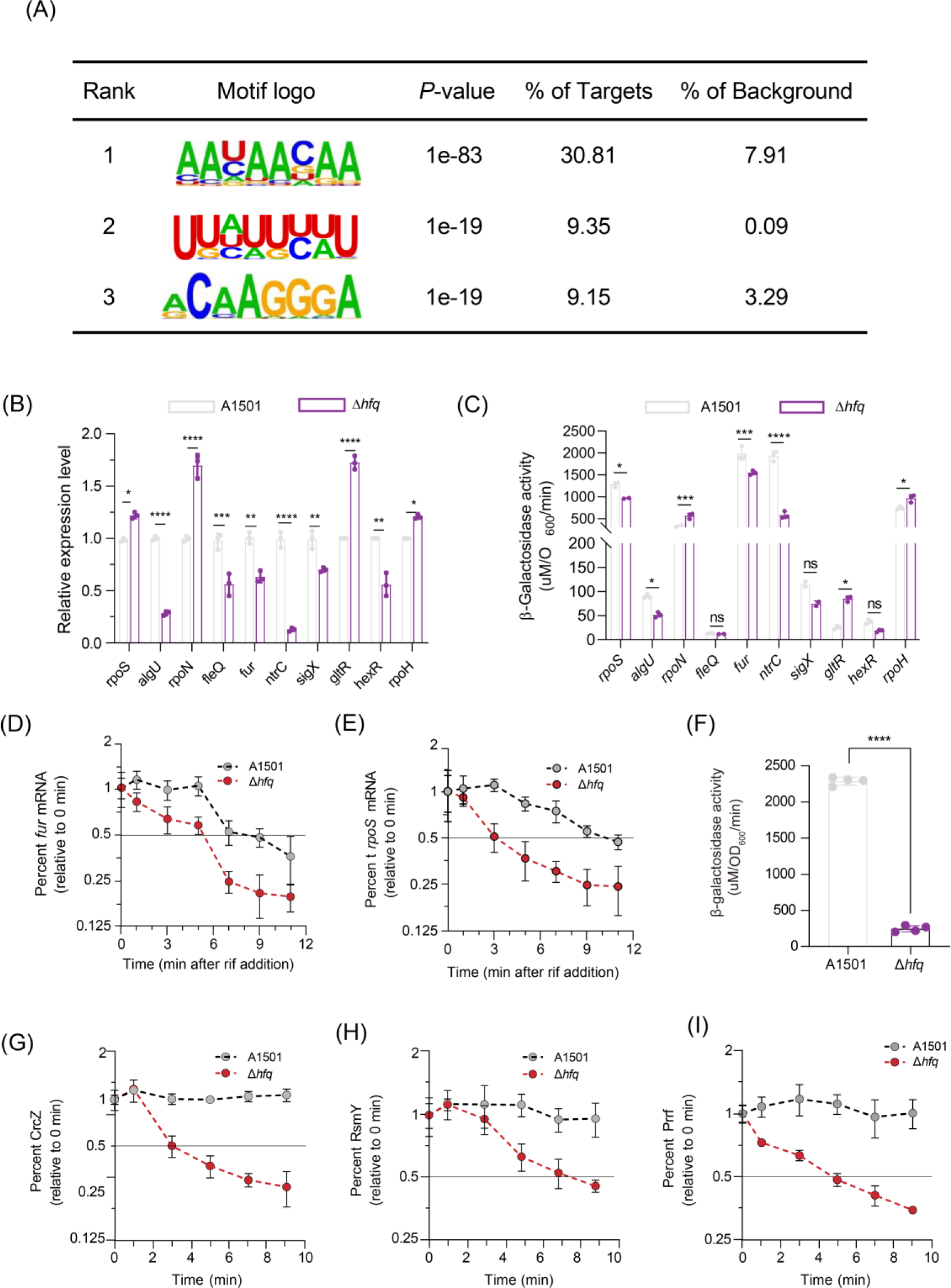
Integration of Hfq into *P. stutzeri* regulatory networks. (A). Top three enriched sequence motifs corresponding to Hfq eCLIP-seq peaks identified with Homer. (B). qRT-PCR analysis of *rpos*, *algU*, *rpoN*, *fleQ*, *fur*, *ntrC*, *sigX*, *gltR*, *hexR*, and *rpoH* RNA levels normalized to the 16S rRNA levels. (C). β-galactosidase activity of the above genes in the WT and Δ*hfq* strains. (D, E). qRT‒PCR analysis of the *fur* (D) and *rpoS* (E) mRNA half-life in the WT and *hfq* mutant strains. Rifampicin (400 μg/mL) was added at time 0. The error bars show the standard deviations of the means of three independent experiments. (F). β-Galactosidase levels conferred by the chromosomally integrated transcription P*crcZ*::*lacZ* fusion in the WT and Δ*hfq* strains. (G, H, I). Half-life of CrcZ (G), RsmY (H), and PrrF (I) in the WT and Δ*hfq* strains. The asterisks indicate that a Hfq eCLIP-seq peak is associated with the gene. Asterisks indicate statistical significance determined by a t test (f) and two-way ANOVA (b, c) with Tukey’s post hoc test: \*\*\*\**p* ≤0.0001, \*\*\**p* ≤0.001, \*\**p* ≤0.01, \**p*≤ 0.05, and ns: not significant.

Furthermore, RNA-seq analysis of ncRNAs in the Δ*hfq* and WT strains showed that the knockout of Hfq decreased the abundance of nearly all identified ncRNAs (Table S3). We previously observed that the CrcZ and RsmY ncRNAs antagonize Hfq- or RsmA-mediated translational repression underlying hierarchical carbon substrate utilization and nitrogen-fixing biofilm formation, respectively (9, 28). In this study, the expression and half-life of the both ncRNAs were decreased in Δ*hfq* (Fig. 4G-H). The substantial reduction in CrcZ levels in the absence of Hfq might nonetheless be due not only to diminish P*crcZ::lacZ* activity but also to reduced ncRNA stability, as CrcZ features several A-rich motifs strikingly similar to the RNase E target (Fig. 4F, S4A). Although no eCLIP-seq peak was found to be associated with PrrF, a ncRNA subject to positive regulation by Fur (46), we observed that the half-life of PrrF was only 4.5 min in Δ*hfq*, revealing marked instability in the absence of Hfq (Fug 4I).

We analyzed the distribution of the top three motifs that corresponded to the peaks and harbored strong Hfq-binding sites by using Homer software (47). We identified an “AANAANAA” motif highly enriched in the Hfq eCLIP-seq peaks. This motif had the greatest peak densities in the 5’ UTRs of mRNAs, accounting for 30.8% of all the peaks (Fig. 4A). An analysis of Hfq 3’UTR peaks revealed a motif with a U-rich sequence “UUAUUUUU” (Fig. 4A), which strongly resembles a Rho-independent terminator ending in long single-stranded poly (U) tails, suggesting that Hfq binds to the mRNA 3’ end and influences the expression of transcripts at the posttranscriptional level. We also found strong enrichment of the “GGA” motif of mRNAs, accounting for 9.2% of the peaks (Fig. 4A), which is similar to findings corresponding to the binding site of RsmA, a member of the CsrA family of proteins that bind RNA as a dimer and recognize sites with the consensus sequence GGA sites.

We further observed that Hfq binds to *rsmA* mRNA and represses the translation of *rsmA*, as determined by β-galactosidase activity analysis (Fig. 3G). As well-studied globally acting RNA-binding proteins, Hfq and RsmA are both involved in optimal nitrogenase activity and efficient root colonization. However, their regulatory functions differ, with Hfq exerting positive regulation(9) and RsmA engaging in negative regulation (28). Notably, both proteins recognize the consensus sequence GGA motif, suggesting that Hfq-bound mRNAs might interact with RsmA, indicating potential overlaps, complementary actions, or competitive roles between these two proteins. Overall, this study expands our knowledge of the putative direct regulatory targets of Hfq, offering a foundational resource for guiding future investigations into the intricate regulatory network underlying nitrogen fixation in root-associated bacteria.

## DISCUSSION

Biological nitrogen fixation is an energy-expensive and highly regulated process(48). It is particularly noteworthy that efficient nitrogenase activity requires maintaining sufficient levels of *nif* mRNAs. Indeed, nitrogen-fixing cells have evolved complex posttranscriptional regulatory networks, which include but are not limited to various RNA chaperones and associated regulatory ncRNAs, to produce *nif* mRNAs at a level sufficient to sustain maximal nitrogenase activity (27, 32). By comparison, our understanding of posttranscriptional regulatory networks underlying nitrogen fixation traits is lagging behind that of transcriptional regulatory systems. In nitrogen-fixing bacteria, Hfq has been implicated in the control of various genes, including those related to optimal nitrogen fixation; however, the substantiating evidence for these links is limited to transcriptomic, phenotypic and genetic studies(9,27,32). Limited information is available regarding the global role of Hfq, especially its target mRNAs. Here, we performed comparative eCLIP-seq analysis under nitrogen fixation conditions, shedding light on *nif*-specific RNA regulation by Hfq. As a result, 987 peaks with significant enrichment in cross-linked samples were identified throughout the *P. stutzeri* A1501 genome. This study underscores the pivotal role of Hfq in nitrogen fixation, as evidenced by the extensive array of regulatory ncRNAs, some of which were previously identified as Hfq targets (27, 49). Furthermore, our observations reveal that Hfq-mediated regulation affects virtually every aspect of A1501 physiology in conjunction with other global regulators under nitrogen fixation conditions.

In *P. aeruginosa*, Hfq was needed for RsmA to bind to *vfr* mRNA, which encodes a transcription factor that controls the expression of biofilm-associated genes (50). Another previous ChIP-seq study indicated that both Hfq and RsmA can bind the nascent transcript of AmrZ, an important global transcription regulator that controls motility, virulence, and biofilm formation in *P. aeruginosa* (51). The comparison of *P. aeruginosa* transcriptomic studies performed for RsmA and Hfq regulons identifies numerous overlaps of genes that were differentially expressed between WT and the respective null mutants(51, 52). Recently, new sequencing methodologies have revealed interplay between multiple RNA-binding proteins and their target RNAs (4). Such regulatory interplay was also observed between Hfq and RsmA in A1501, as shown in Fig. 4A. For example, Hfq and RsmA both recognize the consensus sequence GGA motif, implying a potential overlap of their target genes, such as RsmY. Consequently, the prospect of a coupling regulatory mechanism between Hfq and RsmA arises. Our data substantiate this possibility by showing that Hfq directly binds *rsmA* mRNA and enhances its stability under nitrogen fixation conditions, which to our knowledge has not been reported previously. A distinct feature of Hfq-mediated regulation in *Pseudomonas* is that Hfq inhibits the translation of target transcripts by forming a regulatory complex with the catabolite repression protein Crc (53). Notably, our eCLIP-seq data revealed a significant enrichment of Hfq binding to transcripts that are recognized targets of Crc, including *gltR*, *zwf*, and *benR*(9). A common characteristic among genes regulated by Crc is the presence of a catabolite activity (CA) motif within their 5’UTR. This motif has been reported to be AANAANAA, where N is any nucleotide that is recognized by the distal surface of Hfq (54, 55). Furthermore, our eCLIP-seq and RNA-seq results showed that Hfq is also able to affect transcription directly by coupling with several transcription factors, such as AlgU, RpoS, and Fur, or indirectly modulating the translation of sigma factors, such as RpoN, an alternative sigma factor typically associated with motility, quorum sensing, virulence, stress responses and nitrogen fixation in bacteria (Fig. 5, Fig. S5A). These results indicate that Hfq can interact with diverse classes of global regulators, thereby unveiling a novel protein‒ RNA cross-link between nitrogen fixation and various associated metabolic pathways.

**Fig. 5.**
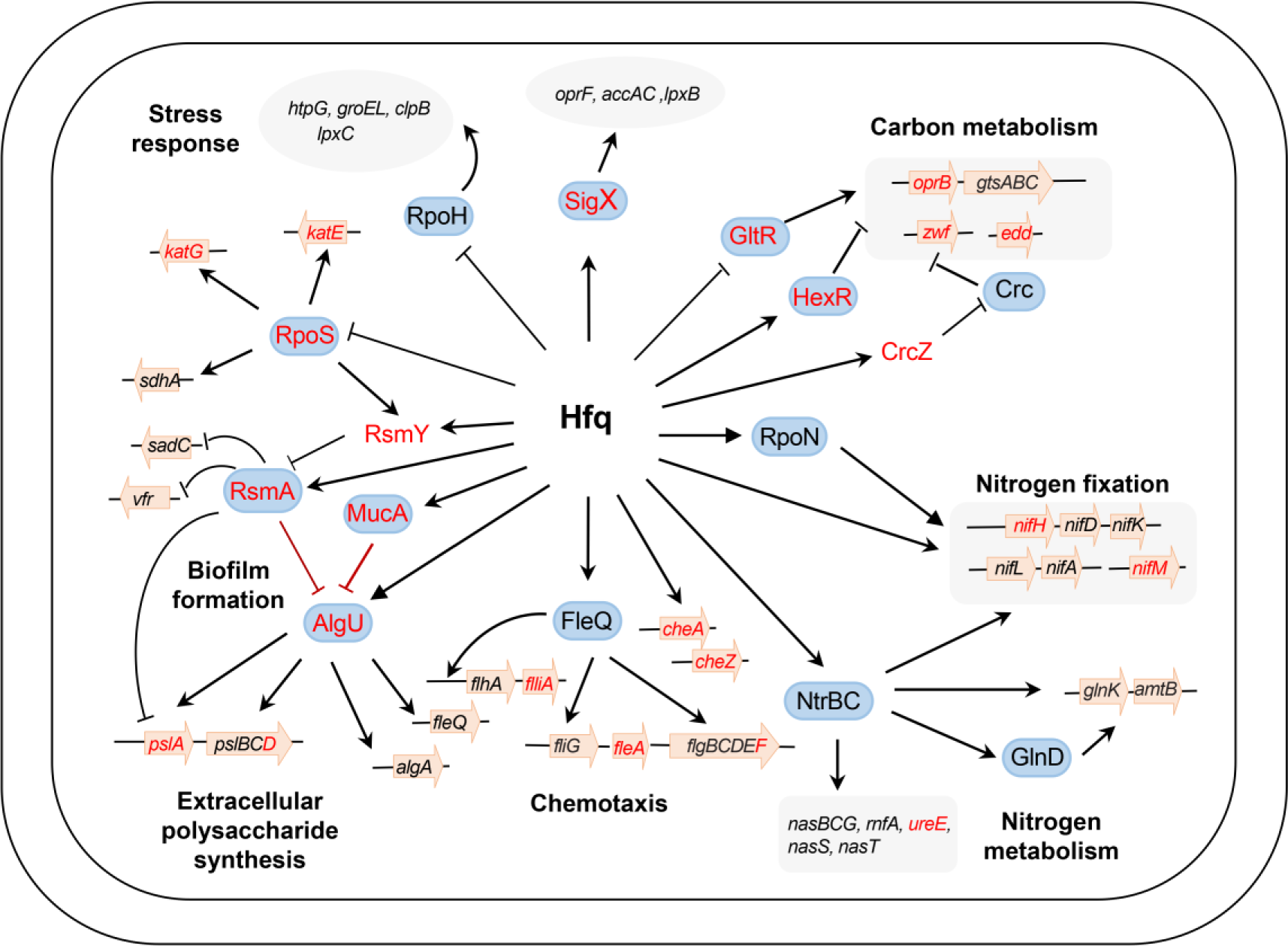
The Proposed regulatory networks controlling nitrogen fixation and other related pathways in *P. stutzeri* A1501. The data are derived from both this study and previous research. In this model, Hfq acts as a central player, working with other global regulators, ncRNAs, and their target genes. Target directly influenced by Hfq are depicted in red, while indirectly affected target are represented in black. Arrows and T-shaped bars indicate positive and negative regulation, respectively. For details, please refer to Table 1 and Supporting Information Table S1.

**Table 1:** Differential expression of selected nitrogen fixation-related genes under Hfq regulation detected by eCLIP-seq and transcriptome studies.

| Gene ID | Gene name | Gene description | *Log<sub>2</sub> Fold Change | P-value |
| --- | --- | --- | --- | --- |
| PST_0503 | *amtB* | ammonium transporter | 0.98 | 3.97E-02 |
| PST_0349 | *ntrC* | nitrogen regulation protein NR(I) | -3.08 | 3.40E-09 |
| PST_0350 | *ntrB* | nitrogen regulation protein NR(II) | -3.09 | 2.31E-06 |
| PST_0353 | *glnA* | glutamate--ammonia ligase Glutamine synthetase, catalytic domain | -0.86 | 2.13E-05 |
| PST_1031 | *rpoN* | RNA polymerase factor sigma-54 | 0.53 | 1.03E-04 |
| PST_1223 | *algU* | RNA polymerase sigma factor | -2.53 | 5.88E-09 |
| PST_1224 | *mucA* | regulatory protein that inhibits *algU* | 1.26 | 2.15E-07 |
| PST_1305 |  | arsenate reductase related protein | 1.94 | 1.42E-04 |
| PST_1306 | *nifB* | GFA family protein | 1.87 | 1.95E-05 |
| PST_1313 | *nifA* | nif-specific transcriptional activator NifA | -1.24 | 9.91E-03 |
| PST_1314 | *nifL* | nitrogen fixation negative regulator NifL | -1.63 | 3.51E-04 |
| PST_1315 | *rnfA* | electron transport complex subunit A | 3.02 | 1.59E-07 |
| PST_1316 | *rnfB* | electron transport complex subunit B | 2.94 | 2.30E-07 |
| PST_1317 | *rnfC* | electron transport complex subunit C | 3.35 | 4.56E-11 |
| PST_1318 | *rnfD* | electron transport complex subunit D | 2.98 | 3.56E-08 |
| PST_1319 | *rnfG* | electron transport complex subunit G | 1.82 | 4.45E-04 |
| PST_1320 | *rnfE* | electron transport complex subunit E | 1.50 | 2.36E-03 |
| PST_1321 | *rnfH* | RnfH family protein | 1.47 | 5.70E-03 |
| PST_1322 | *nifY2* | dinitrogenase iron-molybdenum cofactor biosynthesis | 1.67 | 3.00E-04 |
| PST_1324 |  | conserved hypothetical protein | 2.79 | 1.47E-08 |
| PST_1325 |  | SIR2 family protein | 2.46 | 2.75E-07 |
| PST_1326 | *nifH* | Fe protein, nitrogenase reductase | -1.66 | 1.99E-02 |
| PST_1327 | *nifD* | nitrogenase molybdenum-iron protein alpha chain | -1.05 | 1.14E-02 |
| PST_1328 | *nifK* | nitrogenase molybdenum-iron protein subunit beta | -1.90 | 7.30E-06 |
| PST_1333 | *nifE* | nitrogenase iron-molybdenum cofactor biosynthesis protein | 2.03 | 4.55E-06 |
| PST_1334 | *nifN* | nitrogenase iron-molybdenum cofactor biosynthesis protein | 2.25 | 1.90E-06 |
| PST_1335 | *nifX* | dinitrogenase iron-molybdenum cofactor | 2.34 | 2.13E-06 |
| PST_1336 |  | hypothetical protein | 1.71 | 2.35E-04 |
| PST_1337 |  | hypothetical protein | 0.80 | 1.10E-02 |
| PST_1338 | *fdxB* | ferredoxin, 4Fe-4S | 1.32 | 1.01E-02 |
| PST_1339 |  | ferredoxin, 2Fe-2S | 0.92 | 1.19E-02 |
| PST_1340 |  | hypothetical protein | 0.42 | 8.16E-03 |
| PST_1344 | *nifN* | Nif11-like leader peptide family natural product precursor | 0.15 | 1.16E-05 |
| PST_1345 | *modC* | molybdenum ABC transporter ATP-binding protein | 1.02 | 6.51E-05 |
| PST_1346 | *modB* | molybdate ABC transporter | 5.74 | 1.01E-02 |
| PST_1347 | *modA* | molybdenum ABC transporter | 7.52 | 1.64E-04 |
| PST_1349 | *hesB* | iron-sulfur cluster assembly accessory protein | 1.58 | 7.58E-03 |
| PST_1350 | *nifU* | Fe-S cluster assembly protein NifU | 1.92 | 3.53E-04 |
| PST_1351 | *nifS* | cysteine desulfurase | 1.23 | 1.04E-05 |
| PST_1352 | *nifV* | homocitrate synthase | 1.58 | 3.95E-03 |
| PST_1355 | *nifW* | nitrogenase-stabilizing/protective protein NifW | 0.36 | 2.52E-04 |
| PST_1356 | *nifZ* | Fe-S cofactor synthesis protein | 0.69 | 8.19E-07 |
| PST_1357 | *nifM* | putative a peptidyl-prolyl cis/trans isomerase | 0.02 | 1.77E-04 |
| PST_1359 | *nifF* | flavodoxin required for electron transfer to the Fe protein | -0.23 | 5.12E-04 |
| PST_1371 | *rsmA* | carbon storage regulator | -5.93 | 1.47E-31 |
| PST_1386 | *cheV* | chemotaxis protein | -1.68 | 8.33E-05 |
| PST_1572 | *rpoS* | RNA polymerase sigma factor | 1.53 | 2.40E-03 |
| PST_1993 | *glnT* | type III glutamate--ammonia ligase | 1.35 | 1.53E-02 |
| PST_2442 | *gltR* | the transcription factor required for glucose transport | 6.40 | 3.49E-03 |
| PST_2329 | *zwf* | glucose-6-phosphate dehydrogenase | -1.37 | 8.36E-03 |
| PST_2436 | *oprB* | carbohydrate porin | 1.79 | 3.58E-03 |
| PST_2501 | *pslD* | polysaccharide biosynthesis/export family protein | -2.88 | 5.86E-11 |
| PST_2563 | *motD* | flagellar motor protein | -2.53 | 1.32E-06 |
| PST_2567 | *cheZ* | protein phosphatase | -1.40 | 1.72E-03 |
| PST_3293-PST_3294 | CrcZ | noncoding regulatory RNA | -10.50 | 6.09E-64 |
| PST_3330 | *fur* | ferric iron uptake transcriptional regulator | -1.73 | 5.29E-05 |
| PST_3857-PST_3858 | RsmY | noncoding regulatory RNA | -7.94 | 3.25E-43 |
| PST_3989 | *rpoH* | RNA polymerase sigma factor RpoH | 1.99 | 8.99E-05 |

\*Log<sub>2</sub> Fold Change: The fold change of gene expression in $\Delta hfq$ compared with wild type grown under nitrogen fixation conditions;
Genes enriched in Hfq eCLIP-seq peaks are shown in red;
Genes within the nitrogen fixation island are shaded in gray.
Only those genes referred to explicitly in the text are listed here. For a complete list, see Supporting Information Table S1.

Hfq is widely recognized as a global regulator of cell physiology, and the absence of Hfq results in pleiotropic phenotypic alterations that compromise the fitness of the bacteria and responses against stressful environmental conditions, which particularly affects colonization of the host (56, 57). In *Pseudomonas*, Hfq is important for niche adaptation, and the deletion mutants displayed strongly reduced motility, decreased exopolysaccharides and severely compromised rhizosphere colonization factors (40). Microarray analysis showed that Hfq attenuates the stability of mRNA, explaining the differential expression of ABC genes in *Rhizobium leguminosarum* and *Pseudomonas aeruginosa*(58, 59). Furthermore, transcriptomic and phenotypic data showed Hfq repression of both solute-binding proteins and amino acid ABC transporters in *S. meliloti* (24). Indeed, the deletion of *hfq* resulted in phenotypes by decreased exopolysaccharide production, impaired biofilm formation, compromised motility, and hindered rhizosphere colonization, as revealed in A1501 (9). Δ*hfq* also displayed a significant increase in the abundance of proteins associated with ABC transport and genes related to carbon and nitrogen metabolism (Fig. 5, Table S1). Numerous mRNAs involved in amino acid, nucleotide, and carbon substrate metabolism showed differential upregulation or downregulation in Δ*hfq.* This suggests that Hfq may represent an adaptive response to fluctuations in carbon/nitrogen availability induced by elevated ABC transporter levels. The multifaceted effects of Hfq, as observed in this study, underscore its role as a global regulator governing carbon metabolism, nitrogen fixation, motility, biofilm formation, and fitness, implying a key role in rhizosphere colonization. However, its target genes and the regulatory mechanisms involved in rhizosphere colonization remain unknown.

Nitrogen-fixing bacteria interacting with host plants hold significant agricultural importance. *P. stutzeri* A1501 has emerged as a model organism for investigating the global regulation of nitrogen fixation and microbe-host interactions. Here, we used eCLIP-seq to identify hundreds of novel Hfq-binding RNAs that are predicted to be involved in metabolism, environmental adaptation, and nitrogen fixation. A puzzling aspect of the study is that 121 novel Hfq-binding RNAs are functionally unknown, some of which are previously undiscovered but highly conserved among nitrogen-fixing bacteria. Although Hfq-mediated networks are one of the most extensively studied regulatory networks in bacteria, these functionally uncharacterized RNAs add a layer of complexity to these networks, for example, extending to more connections, such as microbe-host crop interactions. Further experiments will be needed to understand the physiological roles and underlying molecular mechanisms of these RNAs and to elucidate why this diazotrophozth exhibits such remarkable adaptability in rhizosphere environments.

## METHODS

### Construction of Flag-tagged plasmids

Using a fusion PCR protocol, oligonucleotides encoding a 3xFlag affinity tag (DYKDHDGDYKDHDIDYKDDDDK) were added before the TGA termination codon of the *hfq* gene to construct the C-terminally Flag-tagged plasmids. Briefly, a fragment covering an upstream 200-bp region, the entire *hfq* gene followed by 3xFlag and 100 bp downstream were recombined into the linearized vector pLAFR3 using a ClonExpress MultiS One Step Cloning Kit (Vazyme), and the resulting vector was designated pL3xFlag-Hfq. The other fragment spanning an upstream 200-bp region, the first 3 bp of the *hfq* gene followed by 3xFlag and 100 bp downstream, was recombined to yield the pLAFR3 vector using a ClonExpress MultiS One Step Cloning Kit (Vazyme), and the resulting vector was designated pL3xFlag.

### Strains and growth conditions

The strains, plasmids, and oligonucleotides used in this study are listed in Tables S4 and S5. The *hfq*-deletion mutant was generated by homologous double crossover as previously described (9). The Δ*hfq* (pL3xFlag-Hfq) and Δ*hfq* (pL3xFlag) strains were constructed by transferring pL3xFlag-Hfq and pL3xFlag to the Δ*hfq* strain, respectively. *P. stutzeri* A1501 and its derived strains were grown on LB media or minimal medium K (containing 0.4 g L^−1^ KH_2_PO_4_, 0.1 g L^−1^ K_2_HPO_4_, 0.1 g L^−1^ NaCl, 0.2 g L^−1^ MgSO_4_·7H_2_O, 0.01 g L^−1^ MnSO_4_·H_2_O, 0.01 g L^−1^ Fe_2_(SO_4_)_3_·H_2_O, and 0.01 g L^−1^ Na_2_MoO_4_·H_2_O; pH 6.8). Growth experiments were conducted using minimal medium K containing NH ^+^ (6 mM) and succinate (20 mM) as nitrogen and carbon sources. All growth experiments were conducted at 30 ℃ under shaking at 220 rpm in a shaker. If needed, *E. coli* and A1501 were grown in the presence of 10 μg mL^−1^ tetracycline (Tc) or chloramphenicol (Cm) and 50 μg mL^−1^ hygromycin (Hyg) or kanamycin (km).

### eCLIP-seq experimental procedures

The eCLIP experiments, including oligonucleotide sequences and catalog numbers for all reagents, were performed using standard operating procedures as previously described (19). Briefly, the Δ*hfq*(pL3xFlag-Hfq) and Δ*hfq*(pL3xFlag) strains grown under nitrogen-fixation conditions (medium K supplemented with 20 mM succinate) were dispersed onto a petri dish, irradiated with 400 mJ/cm^2^ of UV (254 nm) to crosslink the RNA-protein complex, collected by centrifugation for 8 min at 5,000 rpm at 4 ℃, resuspended in 10 mM Tris-HCl (pH 8.0), and lysed in 1 mL of CLIP lysis buffer (50 mM Tris-HCl pH 7.4, 100 mM NaCl, 1% NP-40, 0.1% SDS, 0.5% sodium deoxycholate, and 1:200 protease inhibitor cocktail III). The resulting sample was subjected to limited digestion for 5 min at 37 ℃ in a Thermomixer at 1200 rpm with RNase I (Thermo Fisher, EN0601), immunoprecipitation of Hfq-RNA complexes with an anti-Flag primary antibody using Dynabeads^TM^ Protein A (Thermo Fisher, 10002D) overnight at 4 ℃, and stringent washes with wash buffer (20 mM Tris-HCl pH 7.4, 10 mM MgCl2, and 0.2% Tween-20). After dephosphorylation with FastAP (Thermo Fisher, EF0654) for 15 min at 37°C in a Thermomixer at 1200 rpm, the sample was incubated with T4 PNK (Vazyme, N102) for 20 min at 37°C in a Thermomixer at 1200 rpm, and stringent washes were then performed. A barcoded RNA adapter was ligated to the 3’ end (T4 RNA Ligase, Thermo Fisher, EL0021). Ligations were performed on-bead (to allow washing away unincorporated adapter) in the presence of a high concentration of PEG8000, which improved the ligation efficiency to > 90%. Hfq-RNA complex samples were then run on standard SDS‒ PAGE protein gels and transferred to nitrocellulose membranes, and a region of 15 kDa extending to ∼75 kDa band size was isolated and treated with proteinase K (Trans gene, GE201) digestion buffer for 20 min at 37°C in a thermomixer at 1200 rpm to isolate RNA. After purification, RNA was reverse transcribed with HiScript III Reverse Transcriptase (Vazyme, R302) and treated with VAHTS DNA Clear Bead (Vazyme, N411) to remove excess oligonucleotides. A second DNA adapter (containing a random-mer of 10 (N10) random bases at the 5′ end) was then ligated to the cDNA fragment 3′ end (T4 RNA Ligase, NEB) in the presence of a high concentration of PEG8000 and dimethyl sulfoxide (DMSO). After cleanup (Dynabeads MyOne Silane, Thermo Fisher), the cDNA was PCR amplified and size selected via agarose gel electrophoresis and VAHTS DNA Clean Beads. The samples were sequenced on the Illumina HiSeq 4000 platform to yield two paired-end 55-bp (for N10) reads.

### eCLIP-seq data processing and analysis

A detailed description of the processing pipeline, including the packages used for the basic processing of eCLIP datasets, was previously described (19, 20, 21, 60). Briefly, eCLIP-seq libraries with distinct in-line barcodes were demultiplexed using custom scripts, and the random-mer was appended to the read name for later usage. After estimating the quality of the raw data using FastQC software, Cutadapt (V.3.4) was used to trim the reads, adapters were processed from the 3’ end of the trimmed reads using a tiling strategy that segments the InvRil19 (/5phos/rArGrArUrCrGrGrArArGrArGrCrGrUrCrGrUrG/3SpC3/) adapter and then mapped to the *P. stutzeri* A1501 genome with STAR. Repeat-mapping reads were segregated for separate analysis. PCR duplicate reads were removed, and then the software SAMtools (version 1.11) was used to sort BAM files and create a BAM index for downstream use. Peak calling with the tool Clipper and motif finding for Hfq were performed with Homer2. Peaks with an FDR-adjusted *p* value <0.1 were considered significant and were used for all downstream analyses. A size-matched input (SMInput) sample was used to normalize and calculate the fold-change enrichment within enriched peak regions using custom Perl scripts (https://github.com/YeoLab/gscripts/tree/1.1/perl_scripts).

For all analyses related to annotated genomic features (CDSs, 5’UTR, and 3’UTR), gene annotations from NCBI were used. We defined transcriptional units (TUs) based on NCBI CDS annotations, both 5’UTR annotations and Rho-independent terminator predictions from Holmqvist (22). Briefly, TUs were defined as starting from the annotated primary transcription start site and ending at either a predicted Rho-independent terminator or in the presence of an intergenic gap larger than 500 bp on the coding strand. In the absence of an upstream transcription start site or a downstream terminator, an arbitrary 200 nt 5’UTR was added upstream of the first codon of CDS, and similarly, an arbitrary 200 nt 3’UTR was added downstream of the last codon CDS. We defined the 5’UTR as the region from the start of each predicted TU to the position upstream of the first codon of CDS in the TU and the 3’UTR as the region from 1 nt downstream of the last CDS to the end of the TU. This raw annotation was then subjected to manual checking, leading to the possible curation of predicted UTRs.

### RNA-seq preparation and data analysis

For RNA-seq, total RNA was extracted from the Δ*hfq* and WT strains under conditions consistent with those used for eCLIP-seq using an innuPREP RNA Mini Kit (Analytik Jena) according to the manufacturer’s instructions. The 23S and 16S rRNAs were depleted using a MICROBExpress bacterial mRNA enrichment kit (Thermo Fisher, USA). For high-throughput sequencing, the libraries were prepared following the manufacturer’s instructions and subjected to Illumina sequencing by Novogene Tech (Tianjin, China). Similar to eCLIP-seq data analysis, the reads were adapter-trimmed, and the remaining reads were mapped to the *P. stutzeri* genome using STAR. Differential expression was identified using DEseq2 (with significance of *p* ≤ 0.05 and a log_2_^(Fold Change [FC])^ threshold ≥1.0 or ≤ −1.0).

For all analyses related to annotated potential ncRNAs, Rockhopper was used to identify new intergenic region transcripts, BlastX was used for comparisons with the nr library to annotate the newly predicted transgenic regions, rfam-scan v1.0.2 with the Rfam c10.0 and BSRD databases was used to annotate ncRNAs, and the unmarked transcripts were used as candidate noncoding RNAs. RNAfold (1.8.5) and IntaRNA (1.8.5) were used to predict the secondary structures and target genes, respectively.

### Quantitative real-time RT‒PCR (qRT‒PCR) analysis

To determine the expression of the indicated genes in WT and Δ*hfq*, total RNA was isolated with an innuPREP RNA Mini Kit (Analytik Jena) according to the manufacturer’s instructions. Total RNA was reverse transcribed using random primers and the High Capacity cDNA Transcription Kit (Applied Biosystems) according to the manufacturer’s instructions. PCR was carried out with Power SYBR Green PCR Master Mix on an ABI Prism 7500 Sequence Detection System (Applied Biosystems) according to the manufacturer’s recommendations. The 16S rRNA gene was used as the endogenous reference control, and relative gene expression was determined using the comparative threshold cycle 2−ΔΔCT method. The data were analyzed using ABI PRISM 7500 Sequence Detection System Software (Applied Biosystems). The primers are listed in Table S4.

### Determination of CrcZ, RsmY, PrrF, *algU, rpoS, fur,* and *rsmA* transcript stability

The stability of CrcZ, RsmY, PrrF, *algU*, *rpoS*, *fur*, and *rsmA* in the A1501 and Δ*hfq* strains, which were grown under nitrogen fixation conditions for 6 h followed by the addition of rifampicin (400 μg/mL, final concentration), was determined. Samples were collected at 0, 1, 3, 5, 7, and 9 min after rifampicin addition and mixed with two volumes of RNA Protect (Sigma) at room temperature for 5 min to immediately stabilize the RNA. The samples were then centrifuged for 5 min at 4 ℃ and 12,000 rpm, and the pellets were rapidly frozen in liquid nitrogen and stored at −80 ℃ until ready for use. Total RNA was isolated with an innuPREP RNA Mini Kit (Analytik Jena) according to the manufacturer’s instructions. cDNA was synthesized from total RNA using a First-Strand cDNA Synthesis Kit (Takara Bio) and was used to estimate the mRNA levels by qRT‒PCR. The primers used are listed in Supplemental Table S4. The relative mRNA concentration was calculated by the comparative threshold cycle (2*^−ΔΔCT^*) method with 16S rRNA as the endogenous reference. The data are presented as the percentages of the transcript levels relative to the amount of these transcripts at time point zero.

### Western blotting

Bacterial cells were lysed in ice-cold wash buffer (1x PBS, 0.1% SDS, 0.5% NP-40 and 0.5% sodium deoxycholate) supplemented with a protease inhibitor cocktail (Roche) and incubated on ice for 30 min. The samples were boiled for 10 min in boiling water with 1x SDS sample buffer, separated by SDS-PAGE, and then electroblotted onto 0.2 μm polyvinylidene difluoride membranes with a Criterion blotter (Bio-Rad). The membranes were blocked with 20 mM Tris-buffered saline and 0.1% Tween-20 (TBST) containing 5% nonfat milk for 1 h at room temperature and incubated with anti-Flag antibody for 1 h and then with HRP-conjugated secondary antibody. Bound secondary antibody was detected using enhanced chemiluminescence reagent.

### Construction of *lacZ* fusion and β-galactosidase assays

To construct transcription and translation gene fusions, the DNA fragments that carried the respective promoter of the target gene were amplified using the primers shown in Table S4 and cloned and inserted into the pUC18-mini-Tn7-Gm-*lacZ* and pXY2 plasmids (61), respectively. The recombinant plasmid pUC18-mini-Tn7-Gm-*lacZ* or pXY2 was electroporated into *P. stutzeri* A1501 together with pUX-BF13, and the mini-Tn7 element carrying the *lacZ* reporter fusion was integrated into the unique Tn7 site located downstream of *glmS*. Specific β-galactosidase activity from bacterial suspensions growing in liquid cultures was measured using 4-methylumbelliferyl-b-D-galactoside (4MUG) as the enzymatic substrate. The fluorescent product, 7-hydroxy-4-methylcoumarin (4MU), was detected at 460 nm after excision at 365 nm using a FlexStation3 Plate Reader (Molecular Devices). Enzyme activity is reported as μM·OD_600_ ^−1^·min^−1^.

### Nitrogenase activity assays

The nitrogenase activity of bacteria was evaluated using the acetylene reduction assay according to a previous protocol (62). After overnight culture in LB medium, the cells were centrifuged and resuspended in a 100 mL flask containing 10 mL of minimal medium K supplemented with 20 mM succinate as a carbon source to an OD_600_ of 0.1. The suspension was subsequently incubated for 24 h at 30°C under an argon atmosphere containing 0.5% oxygen, and 10% acetylene was then added. Gas samples (0.25 mL) were collected at 2 h intervals to determine the amount of ethylene produced on a poly divinylbenzene porous bead GDX-502 column using an SP-2100 gas chromatograph fitted with a flame ionization detector (Beijing Beifen-Ruili Analytical Instrument Co., Ltd.). The ethylene content in the gas samples was determined in reference to an ethylene standard. The nitrogenase activity was expressed as nmol ethylene min^−1^ mg^−1^ protein. The protein concentrations were determined using a Bio-Rad protein assay reagent kit (Bradford, Bio-Rad).

### Biofilm formation assays

Surface-adhered biofilm formation was assayed using the crystal violet method with 96-well microtiter plates (28). The strains used for biofilm experiments were grown overnight in LB at 30°C. The cultures were centrifuged and diluted to a final OD_600_ of 0.2 in fresh minimal medium K. Two hundred microliters of each culture was aliquoted into separate wells in 96-well PVC plates. Microtiter plates were placed in a 30°C incubator without agitation for 48 h. Nonadherent planktonic cells were removed using a multichannel pipette without disturbing the biofilm area, and individual wells were washed twice with 160 μL of sterile double-distilled H_2_O. Then, 160 μL of 0.1% CV solution in ethanol was added to each well for 10 min, and the plate was washed four times with 200 μL of ddH_2_O. After a photograph was obtained, the cell-associated CV was solubilized with 30% acetic acid and quantified by measuring the OD_560_ of the resulting solution using a spectrophotometer.

### Chemotaxis and swimming assay

The chemotaxis phenotype was assessed on soft agar plates (2.5 g of agar per liter) containing succinate as the carbon source(63),(64). The A1501 and Δ*hfq* strains were grown overnight in LB at 30 °C, and the cultures were then harvested, washed twice, and diluted to an OD_600_ of 0.2. Then, 100 mm petri plates were filled with 25 mL of K medium, and the cultures were inoculated by stabbing a toothpick into the agar at the center of the plate. The plates were placed in a 30 °C incubator without agitation for 48 h, and photographs were taken.

### Quantification and statistical analysis

Statistical analysis was performed using the R computing environment and GraphPad Prism 9.0. All data are presented as individual values. A two-tailed unpaired Student’s t test using a 95% confidence interval was used to evaluate the difference between two groups. For more than two groups under different conditions, two-way or one-way ANOVA was used. A probability value of *p*≤0.05 was considered to indicate significance. The experiments were repeated independently three times, and the results were consistent. The data are shown as the averages ±SEMs (standard errors of the mean). ****: *p* ≤ 0.0001; ***: *p* ≤ 0.001; **: *p* ≤ 0.01; *: *p* <0.05; ns: not significant.

## ACKNOWLEDGMENTS

We thank DIATRE Biotechnology (Shanghai, China) for computational analysis assistance. This work was supported by grants from the National Key R&D Program of China (2022YFA0912100, 2019YFA0904700) and National Natural Science Foundation of China (31930004, 32150021, 32270067).

## AUTHOR CONTRIBUTIONS

Y.Z., Y.Y., and M.L. conceived the study and secured funding. F.L., M.L., Y.Z., Y.Y., wrote the manuscript. F.L., M.L., Y.Z., and Y.Y., designed the experiments. F.L., W.S., C.Y., Y.H., Y.S. Y.M., and S.J. performed experiments.

